# A stochastic spatial model for heterogeneity in cancer growth

**DOI:** 10.1101/584573

**Authors:** Alexandre Sarmento Queiroga, Mauro César Cafundó Morais, Tharcisio Citrangulo Tortelli, Roger Chammas, Alexandre Ferreira Ramos

**Affiliations:** Instituto do Câncer do Estado de São Paulo (ICESP), Faculdade de Medicina da Universidade de São Paulo (FMUSP), São Paulo, Brazil; Escola de Artes, Ciências e Humanidades (EACH), Universidade de São Paulo (USP), São Paulo, Brazil; Instituto do Cancer do Estado de Sao Paulo

## Abstract

Establishing a quantitative understanding of tumor heterogeneity, a major feature arising from the evolutionary processes taking place within the tumor microenvironment, is an important challenge for cancer biologists. Recently established experimental techniques enabled summarizing the variety of tumor cell phenotypes in proliferative or migratory. In the former, cells mostly proliferate and rarely migrate, while the opposite happens with cells having the latter phenotype, a “go-and-grow” description of heterogeneity. In this manuscript we present a discrete time Markov chain to simulate the spatial evolution of a tumor which heterogeneity is described by cells having those two phenotypes. The cell density curves have two qualitatively distinct temporal regimes, as they recover the Gompertz curve widely used for tumor growth description, and a bi-phasic growth which temporal shape resembles the tumor growth dynamics under influence of immunoediting. We also show how our representation of heterogeneity gives rise to variable spatial patterning even when the tumors have similar size and dynamics.

**Author summary:** We present a spatial stochastic model to represent the growth of a tumor as a structure having cells of two phenotypes: one whose cells have division as their predominant transition, and another where cells are mostly migrating. The migratory phenotype results from a transformation of the proliferative. Our proposition is based on the assumption that a tumor grows initially within a limited region while its cells are capable of acquire nutrients. During that phase, the cancer cells start changing their phenotype because of stress in their microenvironment and exhaustion of nutrients that lead them to become more migratory and capable of generating metastasis. Our model enables us to recover the usual dynamics observed in tumor growth such as a logistic-like curve, called Gompertz model, widely observed, or the bi-phasic growth observed characterized by equilibrium phase interspersed between two growth regimes. Our approach also enable us to understand the internal spatial and temporal structure of the two sub-populations and can be useful to investigate the phenomena underpinning heterogeneous tumor growth, a feature that helps on the design of treatment strategies based on mitigating heterogeneity related drug resistance.

## Introduction

Despite recent advances on characterization of tumor heterogeneity the understanding of how such a variability affects the tumoral spatial dynamics is still in its infancy [1–3]. Modifications of the tumoral microenvironment exerts an evolutionary pressure that gives rise to new tumor cell phenotypes. At a molecular level, the variety of cellular phenotypes observed in tumors [4] can be connected with the unavoidable randomness of the inner cell environment in which biochemical reactants are present in low copy numbers [5]. Indeed, in tumor cells gene expression levels are highly variable [6–8], and induce the development of phenotypes distinctive by their signal response, a feature that reveals cancer treatment resistance and later relapse [3]. The resisting phenotypes might result from either a cell type originally non-sensitive to a given treatment or result from adaptation of sensitive cells that received an insufficient dosage of antineoplastic agents [9–11]. Such a clinical consequence strengthens the necessity of understanding the biology of tumor heterogeneity and its role in tumor development, a complex task to which effective quantitative models [12] constitute an important additional toolbox.

The inherent randomness of intracellular processes leads to the unique dynamics of tumor development in each tissue and individual. That unique dynamics, which we denote as tumor trajectory, is governed by a probability distribution that results from a plethora of biochemical processes happening inside the cell. Although overwhelming, the complexity of carcinogenesis can be resolved by a combined use of experimental and theoretical techniques appropriate to the investigation of specific phenomena satisfying sufficiently stringent criteria. For example, deterministic models have been employed to describe cancer related processes when an average behavior is observed such as the logistic-like tumor growth dynamics [13–16], cancer invasiveness [17–20], or evolutionary carcinogenesis [21–24], while statistics can be useful to quantify the effects of random fluctuations in tumorigenesis [25,26]. However, those techniques are not sufficient to describe the tumor trajectories, or their probabilities of occurrence, and alternative approaches based on the theory of stochastic processes are essential to investigate that class of phenomena.

In this manuscript we present a cellular level stochastic model for tumor growth where phenotypic heterogeneity is represented in terms of cells being proliferative or migratory. The representation of both phenotypes is effective and has no explicit dependence on DNA sequence or other molecular markers. Our approach enables considering either a “go-or-grow” dynamics [27–29] or a non-exclusive behavior, that we denote as “go-and-grow”, where migratory cells show slow growth capacity, and proliferative cells can migrate slowly [30,31]. Mathematical models approaching the go-or-grow dichotomy have been presented previously with mutually exclusive cell phenotype being determined by the internal molecular quantities [32–37]. In our model, we propose a “go-and-grow” process by attributing a low probability of migration to the proliferative cell, and vice-versa, which has the “go-or-grow” regime as a particular case. To give an effective representation of environmental cues, we propose the cell division rate to decay with tumor size while the death and migration rates increase as sigmoidal functions. The spatial dynamics of our model is simulated by means of a discrete-time Markov chain. Our approach recovers the Gompertz-like growth curve for the tumor size and shows the occurrence of distinctive sub-population dynamics for tumors of similar sizes. We also show the conditions for a two-phase tumor growth and characterize the distinct spatial patterns of cellular sub-populations mixing in a tumor.

## Models and Methods

### A spatial stochastic model for tumor growth heterogeneity evolution

We propose an agent based spatial stochastic model for heterogeneity in tumor growth. The tumor heterogeneity is represented by two sub-populations of cells which phenotypes are predominantly proliferative or migratory. The parameters accounting for the proliferative (and migratory) phenotypes will have indices *p* (and *m*). Our phenotypic classification indicates that during a given time interval a proliferative cell has a higher probability of division than of migration while the opposite happens if we have a migratory phenotype. We assume that the tumor starts with a proliferative cell that can become migratory with non-null probability. For simplicity, in this manuscript we assume that the migratory cells do not transform into proliferative. Table 1 summarizes the symbols and their meaning in throughout this manuscript for our approach and for the Gompertz model.

**Table 1.** List of mathematical symbols. We assume an explicit dependence of cell division rate to the population size, and a dependence of the migration and death rates to the division rate.

| Parameters of the stochastic model for tumor heterogeneity |  |
| --- | --- |
| Symbol | Interpretation |
| $i$ | index denoting the cell phenotype, $i = p, m$ |
| $N$ | number of cells inside the grid at time $t$ |
| $\alpha_i(N), \delta_i(\alpha_i), \rho_i(\alpha_i)$ | cell division, migration, death rate of $i$-th phenotype |
| $A_i, D_i, R_i$ | maximal cell division, migration, death rates of $i$-th phenotype |
| $\sigma_i$ | quiescence rate of the $i$-th phenotype |
| $\bar{N}$ | population size for which $\alpha_i(\bar{N}) = A_i/2$ |
| $\bar{\alpha}$ | cell division rates at which $\delta_i/D = \rho_i/R = 1/2$ when $\alpha_i(N) = \bar{\alpha}$ |
| $k$ | slope of transition $\alpha_i(N), \delta_i(N), \rho_i(N)$ with increase in $N$ |
| $\nu$ | rate of transformation of proliferative phenotype into migratory |
| Gompertz model related parameters |  |
| $x(t), K, \gamma$ | cell density, carrying capacity, maximal cell division rate |

In our model, environmental cues are approached effectively by proposing the cell division rate as inversely proportional to the population size. Conversely, the migration and death rates increase as the division rate decreases. There are biological reasons for those assumptions: (1) in *vivo* and *in vitro* tumor cells doubling time increase with tumor growth [13,38,39]; (2) the relative cell growth rates estimated from experimental data decrease over time following both exponential and sigmoid shape [15];(3) migration is a widely conserved evolutionary response of biological systems under environmental resource limitation [11]; (4) there is experimental evidence showing the coexistence of a proliferative and a highly invasive subpopulations with slow division rate in tumor [29].

Our tumor growth model dynamics is constructed considering four transitions of the state of the *i*-th cell phenotype (*i* = *p*, *m*). The *i*-th cell phenotype undergoes division, migration, death, and quiescence, respectively, at rate *α_i_*, *δ_i_*, *ρ_i_*, and *σ_i_*. Our assumption for tumor heterogeneity implies *α_p_* > *α_m_* and *δ_p_* < *δ_m_*. We now set the division rate as a sigmoidal function of *N*, an *ansatz* based on *α_i_* being limited above and below. The maximal rates of division, migration, and death of the *i*-th cell phenotype are, respectively, *A_i_*, *D_i_*, and *R_i_*. Furthermore, we expect a smooth and nonlinear decrease on the rate of cells proliferation as the tumor size increases [40], a condition that effectively indicates the reduction of the resources availability. 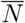 denotes the size of the population at which *α_i_*(*N*) = *A_i_*/2. The migration and death rates are inversely proporional to *α* and reach half of their maximal value when 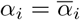. The division, migration, and death rates are written as:

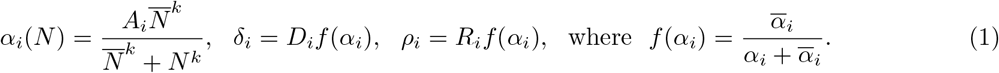

*k* is the slope of the change from maximal to minimal values of *α_i_*(*N*). The tumor size is denoted by *N*, and is measured by the total number of cells at a given time instant, namely *N* = *N_p_* + *N_m_*, with *N_m_* and *N_p_* being the number of migratory and proliferative cells, respectively. In *f*(*α_i_*) the division rate has exponent arbitrarily chosen to be one to avoid introducing more parameters to the model. Fig. 1(A) shows how the cell division rate as function of the total population size. The migration and death rates may be written as functions of *N* if we replace *α_i_* by *α_i_*(*N*) (see Eq. 1):

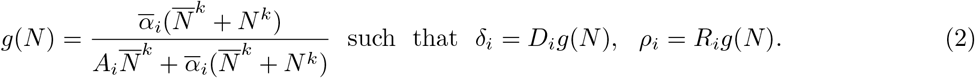

Now, the transition rates of the model are all written as functions of the cell population size (see Fig. 1). In our simulations, we set *ρ_p_* = *ρ_m_* and *σ_p_* = *σ_m_*, and assume that the tumor is in a two dimensional space represented as a square grid of size *n* × *n* – in our simulations we set *n* = 150. Fig. 1(B) shows how the cell migration and death rates depend on the population size.

**Fig 1.**
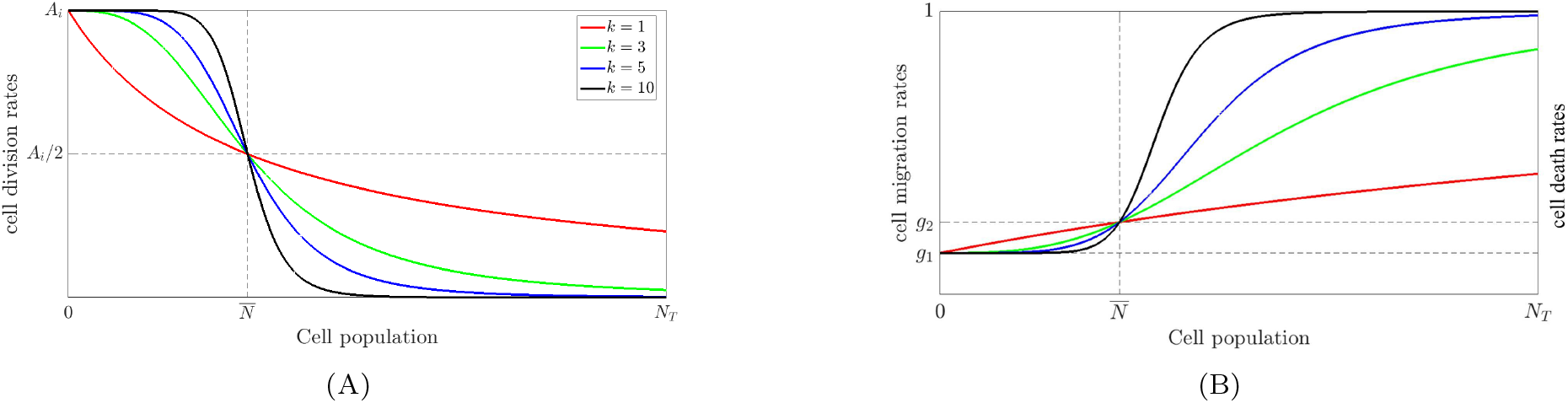
Division, death and migration rates given as functions of the tumor size. *N_T_* is the maximal number of cells accommodated in our grid, namely, *N_T_* = *n* × *n*. (A) The cell division rate as a function of the cell population size (Eq. 1). (B) We plot the Eq. 2 which correspond to the normalized migration and death rates as functions of the tumor size, namely, *g*(*N*) = *δ_i_*(*N*)/*D* = *ρ_i_*(*N*)/*R_i_*. Here 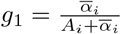 and 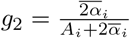.

We denote the probability of a proliferative cell being transformed into a migratory one after division by *P_ν_*, defined as

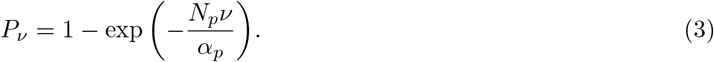

We propose the transformation probability to depend on the division rate of the proliferative phenotype. When the cell division rate decreases there is a higher probability for the phenotype transformation. The use of Eq. 3 for predicting the rate of appearance of the migratory phenotype from the proliferative assumes that a longer doubling-time relates with a reduction on the amounts of metabolic resources. Hence, one may assume a higher probability for the appearance of a migratory phenotype as the value of α reduces (see Eq. 1). Since we are dealing with a pre-invasive regime, here we neglect the phenotypic transition from the migratory to the proliferative state [29]. Note that our formula corresponds to assume an exponential probability of transformation of the proliferative phenotype that is similar to that presumed on the probability of a cancer cell to appear in a tissue after a given amount of cell cycles [41]. Such an *ansatz* is a first approximation and further investigation on it, based on experimental data or new experimental designs, should be encouraged.

### Tumorigenesis dynamics simulation

Fig 2 summarizes our algorithm for simulating emergence of heterogeneity during tumorigenesis. We propose a dynamics based on a finite discrete time Markov chain. At each iteration we select one cell of the population with a probability 1/*N*. Once a cell is selected we consider the transitions that it may perform accordingly with its neighborhood: if the cell has one or more vacant first neighbors the possible transitions are division, migration and death; if there is no vacant first neighbors the possible transitions are quiescence or death. The migratory phenotype is generated with non-null probability during division of the proliferative phenotype. We denote the probability of the *i*-th cell type to perform a transition that has rate *r_i_* (*N*) by *P_r_*(*i*), where the transitions are division, migration, death, quiescence, respectively, denoted by

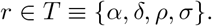

The probability for the transition of rate *r* to happen with the *i*-th cell type is

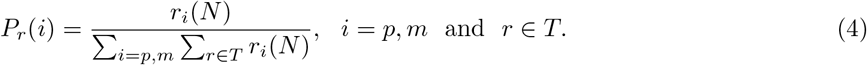

This equation indicates that the probability of a given transition to happen depends on the proportion of its correspondent rate in comparison to the total rate of any transition to happen.

**Fig 2.**
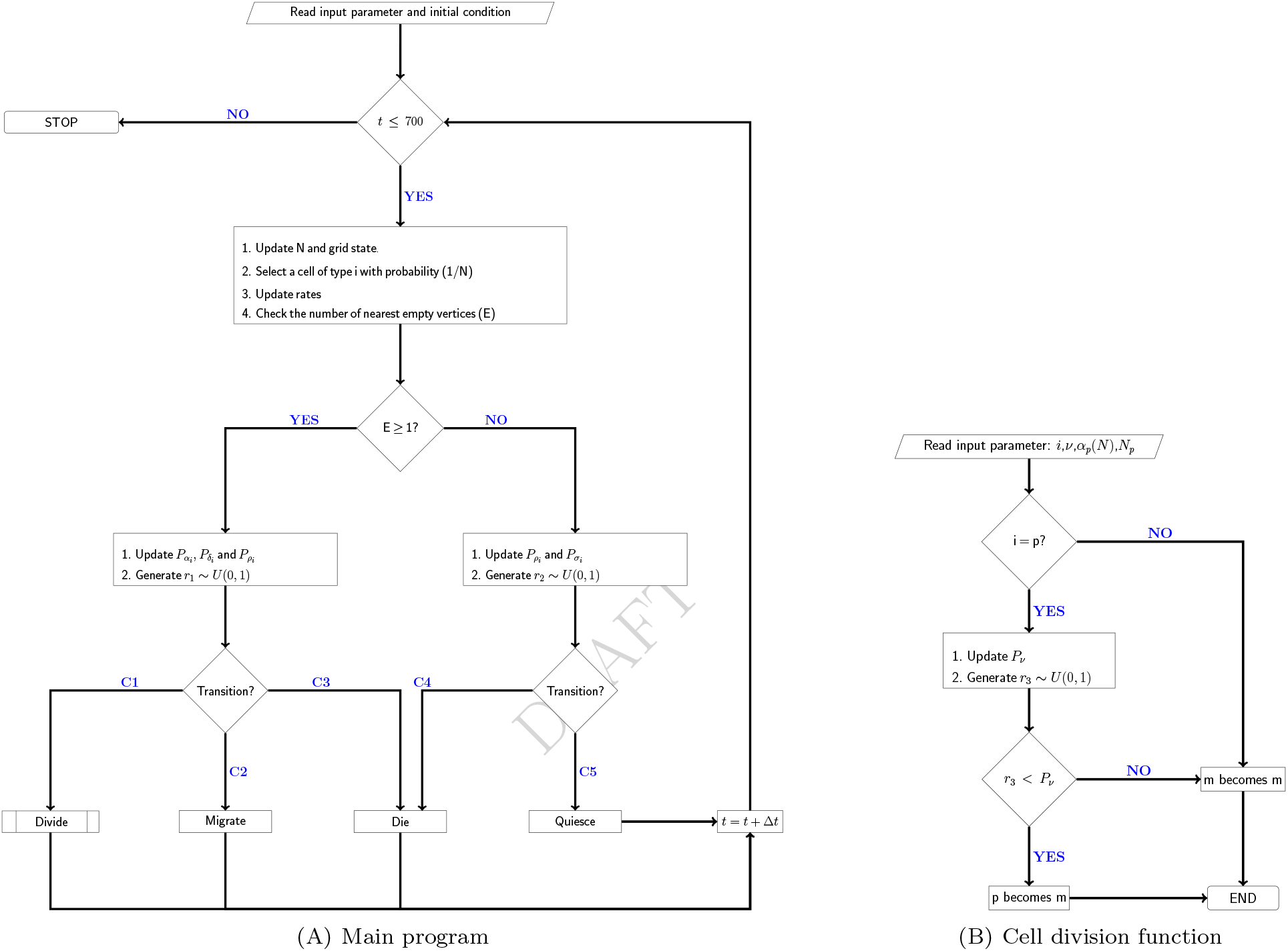
Flowchart representing the algorithm to simulate the tumor growth dynamics. We start with *t* = 0 and at each iteration the time is incremented by Δ*t* as defined by Eq. 5. A single cell is chosen with uniform probability among all cells in the domain. The occupancy of the vertices at one edge distance from the cell is evaluated: *i*) if one or more vertices are empty the cell may duplicate, migrate or die accordingly with probabilities defined by Eq. 4; *ii*) if all vertices are occupied, the cell may quiesce or die. If we chose a proliferative cell that undergoes a division, then there is a probability, given by Eq. 3, that it is transformed in a migratory cell. Let us consider the random numbers *r_i_* and the probabilities of Eq. 4, the probabilistic conditions within the diamonds are given by: (C1) *r*_1_ < *P_αi_*; (C2) *P_αi_* < *r*_1_ < *P_αi_* + *P_δ_i__*; (C3) *r*_1_ > *P_αi_* + *P_δ_i__*; (C4) *r*_2_ < *P_ρ_i__*; (C5) *r*_2_ < *P_σ_i__*. The expression *r* ~ *U*(0,1) indicates a pseudo random number *r* obeying a uniform probability distribution in the interval [0,1).

Although intracellular phenomena are occurring at continuous time our simulations use a discrete time approach and demands the proposition of a correspondence rule between these two time scales. We start assuming that each cell of the tumor is desynchronized from their companions. Thus, one may assume the cell state transitions to be randomly distributed among all *N_i_* cells of *i*-th phenotype of the population. During a given time interval Δ*t* the expected amount of the *i*-th cell type undergoing division (denoted by *L_α_*(*i*)), migration (*L_δ_*(*i*)), death (*L_ρ_*(*i*)), and quiescence (*L_σ_*(*i*)) satisfies: *L_r_*(*i*) ∝ *r_i_N_i_*Δ*t*, with *r* ∈ *T*. For example, the amount of divisions of cell type *p* during the interval Δ*t* satisfies *L_α_*(*p*) ∝ *α_p_N_p_*Δ*t*. One may compute the expected total amount of transitions occurring during an interval Δ*t* by *Q* ∝ ∑_*i*=*p*,*m*_ ∑_*r*∈*T*_ *L_r_*(*i*) such that the time interval Δ*t* corresponding to one iteration in our Markov chain is estimated by

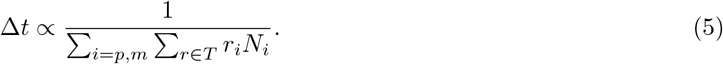

Therefore, at each iteration of our algorithm the time is incremented by the quantity Δ*t* above, and that enable us to relate the discrete time of the simulations with the continuous time of the laboratory.

In our model, the cell density can be defined as the fraction of vertices of the domain occupied by a cell, namely *N*/*n*^2^, *N_p_*/*n*^2^, and *N_m_*/*n*^2^. Our simulations enable us to obtain the dynamics of the cell density within the tumor domain and compare our results with the widely used Gompertz model [13,15]. In the Gompertz model we the population density is denoted by *x*(*t*), the carrying capacity by *K*, and the cell division rate by *γ*. The Gompertz function is written as

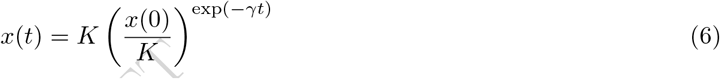

where *x*(0) indicates the initial cell density. The Gompertz function belongs to the class of sigmoidal functions and is bounded above (and below) at *K* (and *x*(0)). At earlier time instants (when *t* << *γ*) it describes a population growing exponentially. For for 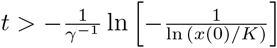 the population density asymptotically approaches *K*, while the growth rate goes to zero. Some examples of the Gompertz curve are shown in blue in Figs.3(A)-3(C).

**Fig 3.**
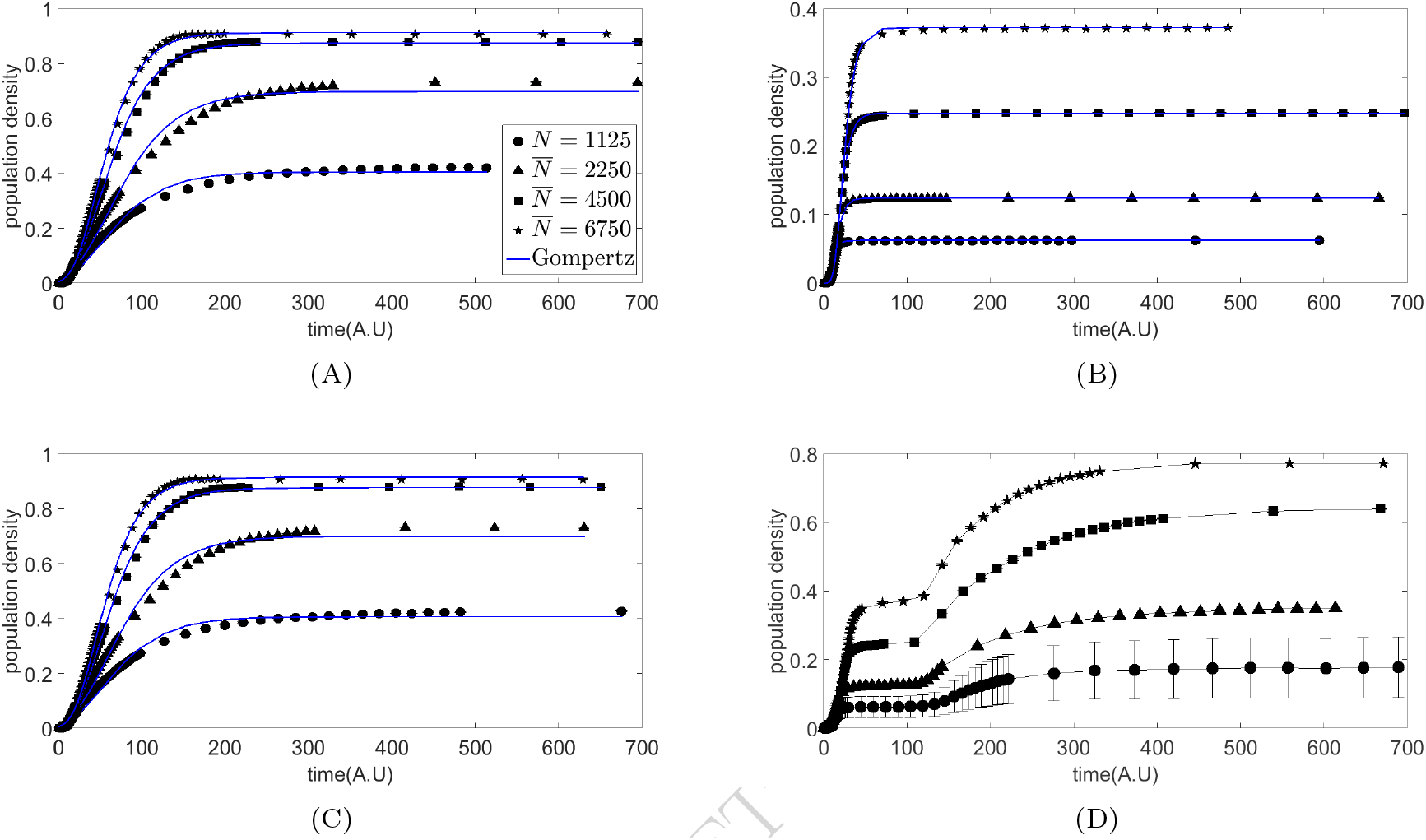
The population density growth accordingly with our simulations. A Gompertz-like growth is shown in Figs. (A)–(C) while a two phase growth is shown in Fig. (D). All trajectories were obtained using 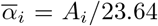, for *i* = *p*,*m*, and *ν* =10^−7^. The symbols related to the values of 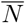 in all trajectories are shown in keys within Fig. (A). The values of *k_i_*’s in Fig. (A) are *k_p_* = *k_m_* = 1, in Fig. (B) are *k_p_* = *k_m_* = 10, in Fig. (C) are *k_p_* = 1, *k_m_* = 10 and in Fig. (D) are *k_p_* = 10, *k_m_* = 1. The parameters for the Gompertz model adjusting to the simulation curves are given in Table 2.

## Results

The simulations of our model are presented considering the dynamics of the total population density, the density of the sub-populations, and the spatial dynamics. We show that our model allows a tumor growth description by a Gompertz-like curve or as a two phases process. Then we show that the proportions of the densities of the two cell phenotypes may differ even in two tumors of the same size at a given time stage. Finally we show how a tumor of the same size can have different spatial distribution of its cell sub-populations. In all simulations we use *A_p_* = 4 *A_m_* = 1, *D_m_* = 10 *D_p_* = 1, *R_m_* = *R_p_* = 1/3, and *σ_p_* = *σ_m_* = 1/100. For each set of parameters used we simulated multiple trajectories to capture stability of our dynamics and its average behavior The time *t* has arbitrary units and frames of the simulation are taken while *t* ≤ 700AU. That corresponds to the steady state of our model, when the population densities stop changing with time. We assume the that the size of the boundaries of the spatial domain of our simulation are fixed and that cells are forbidden to cross them.

### The population density dynamics: Gompertz-like and two phases growth

Fig. 3 shows the population density dynamics obtained by computational simulations of our model for different parameter regimes. We only show the total cell population density for *ν* = 10^−7^. The trajectories for 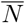 equals to 1125, 2250, 4500, and 6750, are, respectively, indicated by circles, triangles, squares, and stars, as shown in Fig. 3(A) legends. The trajectories at Fig. 3(A), 3(B) and 3(C) are adjusted by a Gompertz function (blue lines), and the fitting parameters given in Table 2 of supplementary material. The curves of Fig. 3(D) show existence of two growth phases followed by a plateau and the black lines in 3(D) are interpolations aiming to simplify the visualization of our results.

**Table 2.** Parameters values for adjusting the Gompertz model to simulation curves shown in Figs. 3(A), 3(B), and 3(C). The Pearson’s correlation coefficient between simulation data and the Gompertz curve is denoted by *r*.

| Figure | $\bar{N}$ | Gompertz-model parameters | | | $r$ |
| --- | --- | --- | --- | --- | --- |
| | | $x(0)$ | $K$ | $\gamma$ | |
| 3a | 1125 | 0.010 | 0.404 | 0.024 | 0.996 |
|  | 2250 | 0.013 | 0.698 | 0.022 | 0.997 |
|  | 4500 | 0.009 | 0.875 | 0.029 | 0.999 |
|  | 6750 | 0.005 | 0.913 | 0.035 | 0.998 |
| 3b | 1125 | 5e-8 | 0.062 | 0.252 | 0.999 |
|  | 2250 | 3e-6 | 0.123 | 0.170 | 0.999 |
|  | 4500 | 5e-6 | 0.247 | 0.129 | 0.999 |
|  | 6750 | 9e-6 | 0.372 | 0.109 | 0.999 |
| 3c | 1125 | 0.010 | 0.406 | 0.024 | 0.991 |
|  | 2250 | 0.013 | 0.699 | 0.023 | 0.992 |
|  | 4500 | 0.009 | 0.877 | 0.029 | 0.995 |
|  | 6750 | 0.054 | 0.914 | 0.035 | 0.997 |

In our simulations all trajectories of total cell population density reach a saturation value. As expected, Figs. 3(A), 3(B), 3(C) and 3(D) show that the saturation density increases with the value of 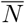 for a fixed set of parameter values. Additionally, Figs. 3(A) and 3(B) indicate that for the same value of 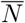 the cell density saturation value is smaller for a higher value of *k*’s. The strength of the regulation of the saturation density can also be noticed on Figs. 3(C) and 3(D) where only one value of *k* is sufficient to induce higher saturation values. Additionally, the smaller value of *k* for the proliferative phenotype is sufficient to ensure this cell to keep dividing even when the population is greater (see Eq. 1) and enables the saturation densities to be comparable to those shown in Fig. 3(A). However, when we have the inverse condition of the *k_m_* being the smaller, there are two growth phases, the former when the population is small and the division rate of the proliferative phenotype is significant, and a second when the population has gone beyond 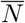 and only the division rate of the migratory population still has a significant value. The Figs. 3(A),3(B) and 3(D) are all adjusted by a Gompertz curve, as generally observed in culture experiments [13,15] while the curve of Fig. 3(D) have also being observed earlier in the context of bacterial growth and were called diauxies [42].

### Analysis of the sub-populations shows diverse dynamics and stationary configurations

Fig. 4 shows the dynamics of the total cell density in black and the corresponding dynamics of the densities of the sub-population of proliferative (in green), or migratory (in red), cells for 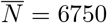. The solid circles (and triangles) are indicating values of the densities when we have *ν* = 10^−5^ (and *ν* = 10^−7^). The parameter values were chosen to demonstrate that at a sufficiently large time the cell density of the whole tumor will be similar. However, Figs. 4(A) – 4(C) show that for the similar stationary cell density one may have different densities of the sub-populations, accordingly with the value of *ν*.

**Fig 4.**
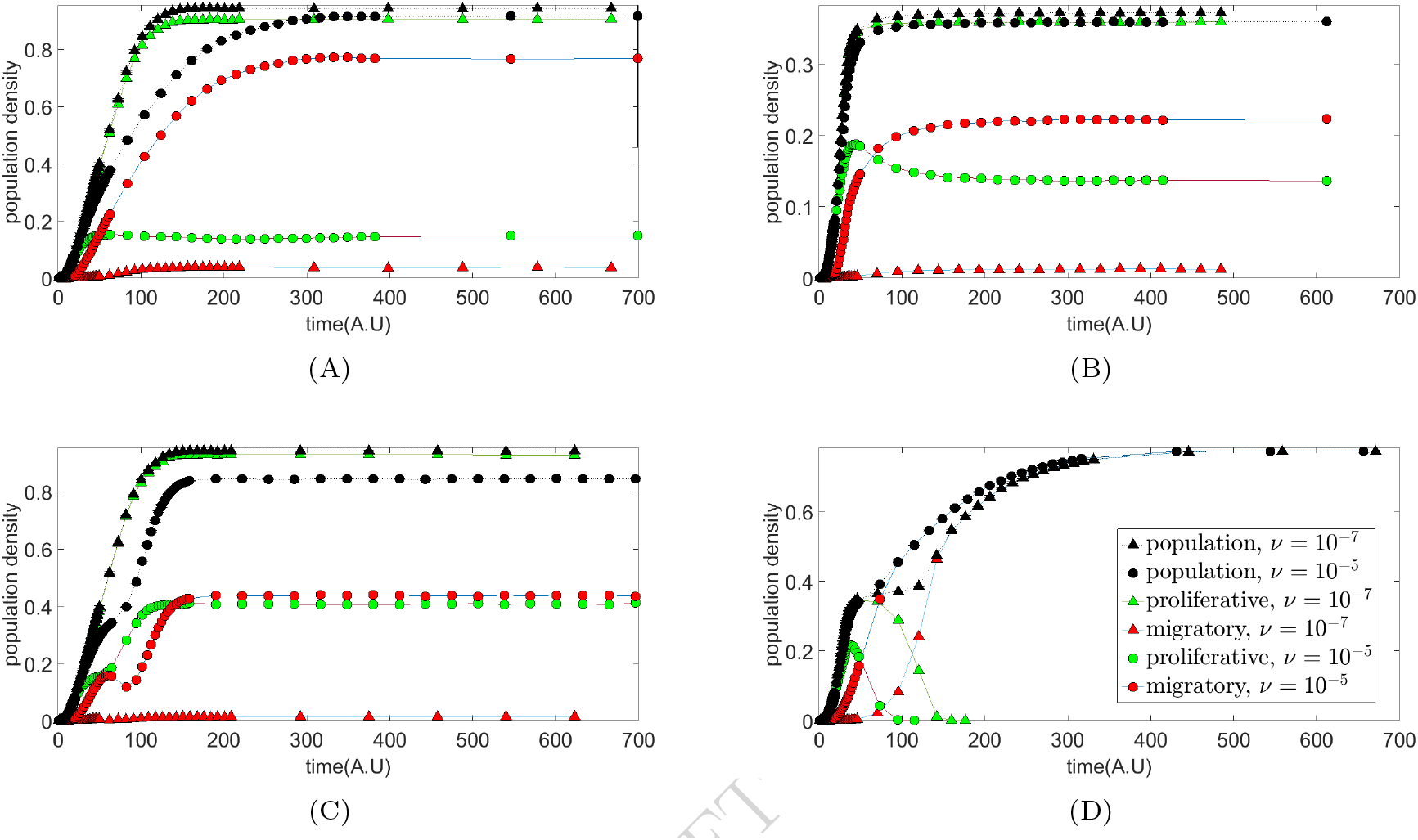
Figures showing sub-populations dynamics underlying total densities growth curve. These four figures brings a summary representing qualitatively different sub-populations evolution over time. For these simulations we set 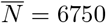. For Figs. (A) and (C) we set 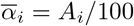, and for Figs. (B) and (D) we set 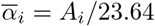. We consider two tumor trajectories, denoted by the solid squares and circles, having *ν* = 10^−7^ and *ν* = 10^−5^. The remaining parameter values are: in Fig. (A) *k_p_* = *k_m_* = 1; in Fig. (B) *k_p_* = *k_m_* = 10; in Fig. (C) *k_p_* = 1, *k_m_* = 10; and in Fig. (D) *k_p_* = 10, *k_m_* = 1.

This is a useful strategy to demonstrate how heterogeneity might develop in a tumor. Fig. 4(A), with *k_m_* = *k_p_* = 1, shows that the speed at which the system reaches the total cell density depends on *ν*. On the other hand, the system reaches steady state at similar speeds in Fig. 4(B) (*k_m_* = *k_p_* = 10). When the values of *k_m_* and *k_p_* are different the dynamics of the total cell density has a stronger relation with *ν*. Fig. 4(C) shows a principle of a diauxie for *ν* = 10^−7^ while there is a clear diauxie for *ν* = 10^−5^ in Fig. 4(D). The sub-populations dynamics also follow different patterns accordingly with the value of *ν*. The growth of the migratory sub-population is faster for *ν* = 10^−5^. During initial instants the proliferative sub-population growth is similar for both values of *ν* but there is always a plateau when *ν* = 10^−5^. Note that the proliferative sub-population is extinguished in Fig. 4(D), and the existence of two non-simultaneous diauxie curves in Fig. 4(C) when *ν* = 10^−5^. Furthermore, in Fig. 4(B) we see that the proliferative sub-population reaches a maximum before reducing towards its steady state density.

### Multiple spatial patterning dynamics gives insights on heterogeneity

Fig 5 presents qualitatively different spatial dynamics obtained with simulations of our model for 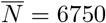, *ν* = 10^−5^. Each row refers to the dynamics obtained with one set of parameters with earlier configurations presented on leftest graphs. All initial conditions are the same: there is one proliferative cell. For 1st row we set *k_p_* = *k_m_* = 1, and 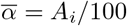 and for 2nd row we set 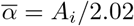.

**Fig 5.**
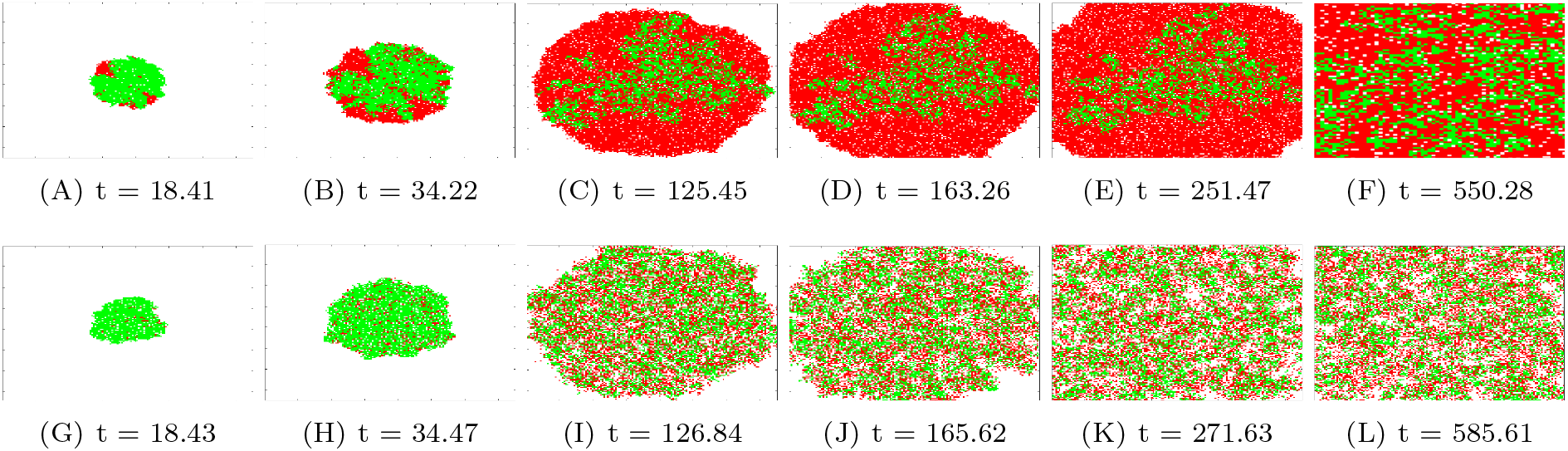
Heatmaps showing spatial configurations of our population dynamics at different time instants. We consider two different qualitative patterns of spatial occupation of the domain. For simulation results shown in both rows we set *k_p_* = *k_m_* = 1, 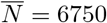, *ν* = 10^−5^ where the results shown at first and second row were, respectively, obtained with 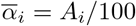 and 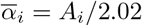.

The heatmaps of the first row, Figs. 5(A),5(B),5(C),5(D),5(E) and 5(F) show that the two sub-populations coexist as two separated phases (approximately), with the migratory population moving to the surroundings and the proliferative remaining at the inner parts of the domain. The second row, Figs. 5(G),5(H),5(I),5(J),5(K) and 5(L) shows a strong mixing between the cells of both sub-populations. In both rows, the population growth has an approximately radial symmetry. The mixing of the sub-populations is understood by means of the higher diffusion rates of the cells in the second row, as we may infer from the values of 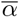 and Eq. 1.

## Discussion

In this manuscript we present a spatial stochastic model for simulating heterogeneity in tumor growth dynamics. We propose an effective approach for describing the cell growth kinetics that recovers the sigmoidal-like dynamics of cell densities. The sigmoidal growth has been widely observed in culture experiments with data being adjusted, for example, by the Gompertz model [15,16,43,44]. The sigmoidal behavior relates with exhaustion of resources in culture experiments which, in our model, is represented effectively by the division rate being dependent of tumor size. Additionally, we also propose that the migration and death rates are inversely proportional to the division rate, to indicate the higher potential of cells to start migrating or dying when resources are scarce. The heterogeneity in our model is represented by means of two cell phenotypes, one being predominantly migratory and the other proliferative. That enables us to investigate how the diverse cellular phenotypes coexisting in a tumor affects its growth. Additionally, it helps us to characterize the variability of spatial distribution of the sub-populations of cells of two tumors of the same size, a potential refinement of cancer staging.

The dichotomic characterization of cancer cells as proliferative or migratory originated in observations made with central nervous system tumor cells line [45], and lead to the formulation of the “go or grow” hypothesis. That implies on considering the migratory and proliferative phenotypes as mutually exclusive, with the tumor cells deterring proliferation to favor migration. However, a recent study using 35 cell lines of tumors originated in three different tissues demonstrated that this exclusive behavior is not general [30]. Indeed, the authors conclude that the cancer cells that they analyzed do not defer proliferation for migration and these two characteristics of a tumor cell are regulated differently depending on their tissue of origin. Such an observation favors our approach as it permits describing the cell phenotypes as an spectrum ranging from mutually exclusive proliferative or migratory towards different combinations of values of the migration and division rates of a given cell, in short, a “go *and/or* grow” description.

One prominent effect of the tumor heterogeneity represented by the two-phenotypes is the diauxie-like curve governing the total cell density. That occurs when the sub-population of proliferative cells grows to a maximum value and decays while the number of migratory cells keep increasing (see Fig. 4(C)). Such a result resembles the cancer immunoediting process having an initially slow growth, when the immune system eliminates modified cells, followed by an equilibrium, when the immune system prevents the tumor to keep growing, and the escape, when the tumor cells overcome the immune suppression and tumor growth restarts with activation of the migratory processes [46,47].

Our approach also allows investigating the spatial patterns of distribution of the two cell phenotypes accordingly with their kinetic constants. Indeed, in each row of Fig. ?? we take a representative pattern. The first row shows a weak mixing of the two cell sub-populations which contrasts with the highly mixed sub-populations at the second row. Note, however, that the populations on those two row grow at similar rates. In the simulation of the first row we consider a small value of 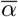, which implies on a slow migration even when we have small values for the cell division rate (see the Eq. 1). Hence, there is a slow spread of cells through the available space and no additional explicit interaction between cells is needed to ensure separation between the sub-populations. On the other hand, the cells of the second column are well mixed and 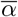 is much greater. That implies on the migration rate that is greater even for smaller division rates such that the two sub-populations become well mixed because of their higher mobility.

Our approach is effective, in the sense that we neglected the microscopic phenomena underlying a given cellular phenotype, such as the switching between the proliferative and migratory phenotypes depending on the amounts of the Mitf and Brn2 [29], or the relation of the cell metabolism and the quantities of RKIP and BACH1 [6]. Establishing the relationship between the cell phenotype and the amounts of its molecular components is an important challenge which would help us to understand the biological meaning of the constant *k*. Fig. 1(A) shows a graph for the division rate as function of *N*, and we note that the greater is *k*, the steeper is the decay of the division rate with the growth of the population. Particularly, note that for *k* = 1 the decay is slow and the division rate may not approach zero. Hence, *k* might also be interpreted as an index for the cell’s sensitivity to contact inhibition [48,49], where lower values of *k* imply on lower sensitivity to contact inhibition, and higher cell densities. Indeed, such an interpretation has some support in our simulations, as we note that lower values of *k*’s result in greater asymptotic cell densities as shown at Figs. 3(A) and 3(B). Additionally, we note that for larger values of 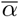, as shown in Fig. 4(D), the sub-population having the smaller *k*, even if it has the migratory phenotype, can reach larger densities.

The occurrence of multiple biological scenarios is expected in a model having multiple parameters and its further refinement will require the guidance of experiments. For example, one might measure the decay of the division rate as function of the cell density to establish a clearer interpretation of the parameter *k*. Additionally, co-culture experiments might be used to understand how *k* affects the prevalence of a given cellular phenotype during the different phases of the tumor growth. Those additional results would support both the verification of the usefulness of our approach and to determine its scope of application in cancer research.

## Acknowledgments

ASQ thanks for the scholarship provided by CAPES and the opportunity to visit Dr. Ariosto Silva at the departament of Integrated Mathematical Oncology at Moffitt Cancer Center-USA. AFR was supported by CAPES (Process n 88881.062174/2014-01, AFR). MCCM thanks CAPES for the scholarship support. Simulations were carried out with High Performance Computing resources provided by the Computer Science Superintendence of the University of São Paulo.

## Supporting information

**S1 Video** Video with frames from simulation which input parameter set were *k_p_* = *k_m_* = 1, 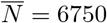, 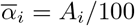, *ν* = 10^−5^. In this simulation we observed that both subpopulation are quite immiscible.

**S2 Video** Video with frames from simulation which input parameter set were *k_p_* = *k_m_* = 1,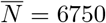, 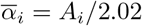, *ν* = 10^−5^. In this simulation we observed that both subpopulation are totaly miscible.

**S3 Video** Video with frames from simulation which input parameter set were *k_p_* = *k_m_* = 10, 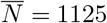, 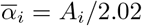, *ν* = 10^−5^. In this simulation we observed that from a given moment the cells spread through the domain.

**S4 Video** Video with frames from simulation which input parameter set were *k_p_* = *k_m_* = 10, 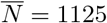, 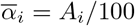, *ν* = 10^−5^. In this simulation we observed that from a given moment the cells remain quite closely.

**S5 Video** Video with frames from simulation which input parameter set were *k_p_* = 10, *k_m_* = 1,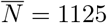, 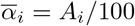, *ν* = 10^−7^. In this simulation we observed that the proliferative cell growth and goes to extinction meanwhile the migratory cells emerge from borders forming cell conglomerates, but in long time intervals the population assume an irregular shape.

**S6 Video** Video with frames from simulation which input parameter set were *k_p_* = 10, *k_m_* = 1,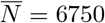, 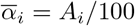, *ν* = 10^−7^. In this simulation we observed that the proliferative cell growth but takes a long time interval to goes to extinction, the migratory cells once again emerge from borders but in this situation the population no longer assumes an irregular shape.

**S7 Video** Video with frames from simulation which input parameter set were *k_p_* = 1, *k_m_* = 10, 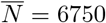, 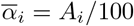, *ν* = 10^−7^. In this simulation we observed that migratory do not last time enough to survive and proliferate.

**S1 Table.** Table containg Gompertz model’s parameters value

